# Fieldable isothermal nucleic acid test for rapid semi-quantitative visual readout of *Enterococci* in recreational waters

**DOI:** 10.1101/2025.04.20.649722

**Authors:** Anirudh Sudarshan, Meredith G. Rillera, Nicholas Tran, Timothy E. Riedel, Andrew D. Ellington, Sanchita Bhadra

**Affiliations:** Department of Molecular Biosciences, College of Natural Sciences, The University of Texas at Austin, Austin TX 78712 USA; Freshman Research Initiative, College of Natural Sciences, The University of Texas at Austin, Austin, TX 78712, USA

## Abstract

*Enterococci* are fecal indicator bacteria whose presence in water suggests the potential for gastrointestinal pathogens. Culture-based detection is slow and may require expensive apparatus and proprietary media. Molecular tests based on qPCR can be technically difficult and require complex devices. Loop-mediated isothermal amplification (LAMP) tests that need minimal instrumentation have been described but they only qualitatively indicate presence or absence of *Enterococci*, which is difficult to compare with quantitative contamination standards for assessing water quality. Here, we demonstrate semi-quantitative ‘thresholded’ LAMP that uses internal competition with pre-defined false targets and an oligonucleotide strand displacement (OSD) probe to measure the degree of *Enterococcus* contamination in water. The assay needs only one hour of incubation at a single temperature following which a simple visual examination of endpoint OSD fluorescence allows order of magnitude scale estimation of *Enterococcus* amounts. Environmental water samples with low, medium, and high *Enterococcus* contamination could be readily distinguished by thresholded LAMP-OSD without interference from non-specific signals. LAMP-OSD performance correlated with the outcomes of both qPCR and plate cultures of the samples. The simplicity of implementing thresholded LAMP-OSD makes it well suited for point-of-need and low-cost water quality monitoring.

## INTRODUCTION

Exposure to gastrointestinal pathogens when recreating in water can lead to significant illness and financial burden.^1^ Traditionally water quality assessment and standards for recreational waters have been based on Fecal indicator bacteria (FIB) like the fecal coliform, *Escherichia coli*, and enterococci, due to their prevalence in human and animal feces.^2^ Current methods for quantitating FIB typically involve culture-based techniques, such as membrane filtration (MF) and most probable number (MPN) testing.^3^ Culture-based methods, though theoretically straightforward and cost-effective, require 18–48 hours to yield results and are prone to contamination and both false positive and false negative errors.^3^ Moreover, in practice, both MPN and MF require a high level of expertise for sample preparation and dilution and interpretation of results.^4^ These limitations hamper the use of MPN/MF for field research since at minimum a kitchen level of infrastructure is required. On the other hand, expensive specialized or proprietary enrichment media and labor-saving automated systems for culture-based sample analysis, such as the IDEXX Quanti-Tray, present a significant cost barrier.

Recently PCR and qPCR-based tests have been developed and due to their increased specificity are allowing for microbial source tracking (MST) between animals.^5^ However, PCR based tests are mostly limited to use within a laboratory setting due to the need for DNA extraction, the high cost of reagents,^6^ expertise demands, and the need for complex instrumentation.^7^ This makes regular monitoring not only expensive but also slow. The fastest documented sample to qPCR answer time was 3.5 to 4 hours after receipt of sample.^8^ While this timeframe can allow same day actionable water quality monitoring, it is only practical if the water body is within ∼1 h transport distance from the testing lab and samples can be collected by 7:00 a.m.^8^ Consequently, <1% of beach water microbial testing is performed using qPCR.^9^

Loop-mediated isothermal amplification (LAMP) of nucleic acids offers a promising alternative to PCR, in terms of potential field applicability.^10, 11^ LAMP can be used to directly analyze crudely processed samples without needing laborious nucleic acid purification.^11-14^ Moreover, by using loop-forming primers and a strand-displacing DNA polymerase LAMP can isothermally produce 10^9^ to 10^10^ copies of target nucleic acids, usually within one hour, thus allowing operation with minimal instrumentation.^10^ However, unlike qPCR, LAMP provides qualitative readout of presence or absence of analytes. This can be particularly limiting for water monitoring where quality assessments are based on measuring elevations in FIB levels rather than their presence alone. A second limitation of LAMP is that it often suffers from spurious amplification that can result in false positives with non-sequence specific readout methods.^15^ We previously overcame both these drawbacks by developing oligonucleotide strand displacement (OSD) probes to produce sequence-specific signal from LAMP amplicons.^16^ The simplest OSD comprises a fluorophore-labeled long strand bound to a quencher-labeled short strand whose displacement by a complementary LAMP amplicon results in fluorescence.^16^ OSDs have been previously used to identify amplicons and SNPs,^16, 17^ compute multiplex amplicons into Boolean output,^12, 14, 15^ transduce LAMP to color or glucose,^18^ and most importantly for this study semi-quantify target templates from a single endpoint readout by thresholding OSD signal by using competing OSD-unresponsive false templates.^19^

Using this more versatile LAMP-OSD platform, we previously demonstrated direct analysis of heat-lysed *E. coli* as well as the rapid semi-quantitation of *Bacteroides* HF183 sequences of human fecal origin in sewage-contaminated water.^11^ Another group recently demonstrated a LAMP assay that targets *Enterococcus* 23S rRNA gene to have comparable specificity and sensitivity to the US EPA *Enterococci* qPCR test for surface water analysis.^20, 21^ However, unlike qPCR, this LAMP assay, read using a DNA intercalating dye, cannot gauge the amount of *Enterococci* in a sample. This is a drawback because determinations of recreational water quality for fecal contamination in fresh and marine water rely on quantitative thresholds, such as a geometric mean threshold of 35 colony forming units (CFU)/100 ml for culture-based detection methods and ≥ 1000 calibrator cell equivalents (CCE)/100 mL for *Enterococcus* qPCR.^22, 23^ Consequently, a LAMP test with a simple yes/no answer for the presence or absence of *Enterococci* may not provide sufficient information for effectively measuring water safety. Therefore, we re-engineered the *Enterococcus* LAMP assay for template semi-quantitation on an order of magnitude scale while retaining the simplicity of a single endpoint visual readout with minimal instrumentation for facile portability. Herein we demonstrate design and evaluation of this semiquantitative *Enterococcus* LAMP-OSD assay and its application in analysis of environmental water samples.

## METHODS

### Chemicals and reagents

All chemicals were purchased from Sigma-Aldrich (St. Louis, MO, USA) unless indicated otherwise. All enzymes were purchased from New England Biolabs (NEB, Ipswich, MA, USA). All oligonucleotides and synthetic DNA were purchased from Integrated DNA Technologies (IDT, Coralville, IA, USA) or Twist Biosciences (Quincy, MA, USA) (**Supplementary Table 1**). *Enterococcus faecalis* NCTC 775 strain was purchased from ATCC (Manassas, Virginia, USA) and *E. faecalis* culti loops (Culti-Loops *Enterococcus faecalis* ATCC 19433) were obtained from Fisher Scientific (Waltham, MA, USA).

### *Enterococcus faecalis* culture

*E. faecalis* was grown overnight at 37°C using ATCC Medium 41: Brain Heart Infusion Agar or Broth (ATCC, Washington D.C., USA). Liquid cultures were aerated at 250 rpm. Prior to use in LAMP-OSD analysis, 1:300 sub-cultures of the overnight cultures were grown in fresh ATCC Medium 41 for four hours in a 37°C shaker.

### OSD Probe Design

Using our previously described design rules,^16^ the NUPACK software (Pasadena, CA, USA),^24^ and the Benchling platform (San Francisco, CA, USA), an OSD probe was designed for a previously reported *E. faecalis* 23S rRNA gene-specific LAMP assay.^20^ The hemiduplex OSD consisted of a 40-nt long 5’-end fluorescein-labeled long strand, designed to hybridize to a 37-nt long region in the loop sequence between the B1 and B2c regions of the LAMP amplicon (**Supplementary Figure 1**). The 3’-end OH group of this strand was blocked against extension by appending an inverted dT residue. A 25-nt long portion of the OSD binding region of the LAMP amplicon was also recognized by the Loop B primer. The OSD short strand modified with a 3’-end Iowa Black quencher was designed for complementarity to 29-nt at the 5’-end of the OSD long strand. This ensures greater thermodynamic stability of the OSD probe versus pairing of the OSD short strand with the 25-nt long Loop B primers. The hemiduplex OSD was pre-annealed by mixing 1 μM of the long strand with 5 μM of the short strand in 1X isothermal buffer (NEB: 20 mM Tris-HCl, 10 mM (NH_4_)_2_SO_4_, 50 mM KCl, 2 mM MgSO_4_, 0.1% Tween®20, pH 8.8@25°C) and incubating at 95°C for 5 minutes followed by slow cooling (0.1°C /sec) to room temperature.

### LAMP-OSD assay

LAMP-OSD reactions were set up in 25 μl volume comprised of 1X isothermal buffer supplemented with 1.2 mM deoxyribo-nucleotides, 0.6 M betaine, 2 mM additional MgSO_4_, 2.4 μM each of FIP and BIP, 1.2 μM each of Loop F and Loop B, and 0.6 μM each of F3 and B3 primers. All assays also received 16 units of Bst 2.0 DNA polymerase (NEB) along with 0.2 μM of OSD long strand pre-annealed with a 5-fold excess of the OSD short strand. In some LAMP assays, amplicon accumulation was evaluated by real-time measurement of dye intercalation by replacing OSD reporters in the reactions with 1X EvaGreen (Biotium, Fremont, CA, USA). Thresholded LAMP-OSD reactions also received indicated amounts of ‘false targets’ ranging from zero to the 10^10^ order of magnitude copies per reaction. The false targets were designed as synthetic *E. faecalis* 23S rDNA in which the OSD binding region was scrambled to prevent hybridization of OSD probes (and Loop B primers). The false targets can be amplified by the remaining five *E. faecalis* LAMP primers without generating corresponding OSD signals.

Performances of LAMP-OSD and thresholded LAMP-OSD assays were tested by adding indicated types and amounts of templates per reaction in 2.5 to 5 μl volume. These included either zero to 10^9^ order of magnitude copies/reaction of synthetic 23S DNA, indicated dilutions of lab-cultivated *E. coli* or recombinant *E. coli* expressing *E. faecalis* 23S rDNA fragment, zero to few thousand colony forming units (CFU) of lab-cultivated *E. faecalis*, or 2.5 μl of filtration-enriched environmental water sample.

All LAMP-OSD assays were incubated at 65 °C for 60 minutes and accumulation of OSD fluorescence was either measured in real-time using a Roche LightCycler® 96 real-time PCR machine (Basel, Switzerland) or imaged at endpoint using a ChemiDoc MP imaging system (Hercules, California) or a blue light transilluminator and cellphone camera.

### 23S TaqMan qPCR assay

PCR primers and TaqMan probe described in the US EPA Method 1611 *E. faecalis* 23S qPCR assay: Enterococci in Water by TaqMan® Quantitative Polymerase Chain Reaction (qPCR) Assay were used for qPCR analyses performed using the Luna® Universal probe qPCR or one-step RT-qPCR Master Mix according to the manufacturer’s instructions (NEB).^21^ The qPCR reactions were incubated at 95 °C for 10 min prior to 45 cycles of denaturation at 95°C for 15 seconds and annealing and extension at 60°C for 2 minutes. TaqMan probe fluorescence was measured in the FAM channel during extension.

### Environmental sample collection and analysis

Environmental water samples were collected from the urbanized and degraded Waller Creek.^25, 26^ Samples were collected at the intersection of Dean Keeton St. and San Jacinto Blvd in Austin, Texas, USA (GPS coordinates DD 30.28857, - 97.73382). Prior to sample collection, sampling bottles were acid-washed with 10% HCl to minimize contamination. Prior to sampling from a shaded, flowing section of the creek bottles were rinsed three times with creek water that was shaken and discarded downstream. Samples were transported on ice and processed within 3 hours.

For *Enterococcus* nucleic acid analyses, 400 mL of creek water samples were filtered using a vacuum manifold and MF-Millipore, 0.22μm mixed cellulose esters (MCE) Membrane qPCR filters. Following filtration samples were crudely processed using the Filter, Heat, Spin protocol described in Pham et al. 2024,^27^ In brief, sterile tweezers were used to fold and transfer the filters into 1.5 microcentrifuge tubes. 500 μL of AE buffer (10 mM Tris-Cl, 0.5mM EDTA, pH of 9.0) was added to the tubes to fully submerge the filters. The tubes were then placed in a 95 °C heat block for 10 minutes followed by room temperature centrifugation at 12,000 rpm (9678 RCF) for 2 minutes and vortex mixing for 10 seconds. Some 250 μL of these processed samples in AE buffer were aliquoted into 1.5 tubes and stored at -20 °C until LAMP or qPCR analysis.

For analyzing *Enterococcus* colony forming units (CFU), 5 mL of the water samples were added to 100 mL of 1X phosphate buffered saline (137 mM NaCl, 2.7 mM KCI, 10 mM Na_2_PO_4_, and 1.8 mM KH_2_PO_4_, pH 7.4) and filtered through white gridded 47 mm MCE S-Pak ® membrane filters of 0.45 μm pore size, then washed with approximately 20 ml of fresh PBS. The filters were then transferred to membrane *Enterococcus* Indoxyl-β-D-Glucoside (mEI) Agar plates (Moltox ® Molecular Toxicology, Boone, NC, USA) using sterile tweezers. The plates were inverted and incubated at 41°C ± 0.5 °C for 24 hours ± 2 hours. Subsequently, colonies that produced a blue halo were presumptively identified as Enterococci and their recorded count was used to calculate the CFU/mL of the water samples.

## RESULTS AND DISCUSSION

### Design of a 6-primer LAMP-OSD assay for *Enterococcus*

As a first step to converting the *Enterococcus* LAMP reaction reported by Martzy *et al*.^20^ into a semi-quantitative assay, a fluorophore and quencher labeled hemiduplex OSD probe was designed to bind the amplicon loop sequence between the B1 and B2c primer recognition regions. Since this region of the loop is also occupied by the loop B primer, the OSD long strand was designed in the same antisense orientation as the loop B primer to prevent primer:probe hybridization. Meanwhile, the binding of the complementary OSD short strand with the loop B primer was thermodynamically disfavored over its hybridization to a longer region of the OSD long strand.

To validate OSD performance, duplicate LAMP assays comprised of all six LAMP primers (FIP, BIP, F3, B3, loop B and loop F) were supplemented with either a DNA intercalating dye or the OSD probe. Similar kinetics of real-time fluorescence accumulation were observed for both the DNA binding dye and the OSD probe in reactions containing different amounts of a synthetic *Enterococcus* 23S DNA template (**Supplementary Figure 2**). Both assay formats consistently detected as few as 1000 template copies with no production of non-specific fluorescence in reactions lacking specific templates. These results indicate that OSD probes could be used for *Enterococcus* LAMP readout without compromising the assay time-to-result, detection limit, or specificity previously described by Martzy et al.^20^ In fact, LAMP-OSD assays read with only a single visual observation of endpoint fluorescence demonstrated the same detection accuracy and sensitivity (**Supplementary Figure 3**). Furthermore, the binding of the OSD long strand and the loop B primer to the same region of the LAMP amplicons did not compromise assay performance as opposed to LAMP-OSD reactions lacking the loop B primer or OSD probes (**Supplementary Figures 2 and 4**).

To verify assay performance with bacterial nucleic acids, LAMP-OSD reactions were seeded with different colony forming units of lab-cultivated logarithm phase *Enterococcus faecalis*. Bright OSD fluorescence was observed at amplification endpoint in LAMP reactions containing as few as 5 CFU of *E. faecalis* (**Figure 1**). As expected, LAMP-OSD assays provided with *E. coli* instead of *E. faecalis* remained as dark as the assays lacking any templates (**Supplementary Figure 5**). These results demonstrate that the 23S rRNA LAMP-OSD assay can be used to detect *Enterococcus* with a single endpoint visual observation of OSD signal without interference from non-specific noise.

**Figure 1.**
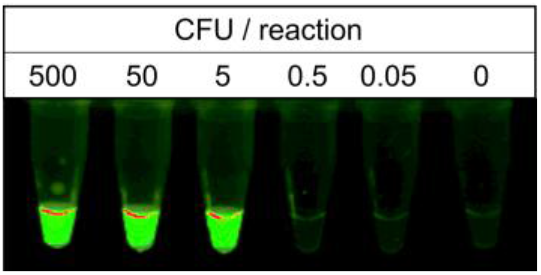
Visual LAMP-OSD analysis of lab-cultivated *Entero-coccus faecalis* bacteria. Image of endpoint OSD fluorescence in LAMP-OSD assays containing indicated colony forming units of log-phase *E. faecalis* is depicted. Data shown is representative of triplicate analysis.

### Development of signal thresholded LAMP-OSD for *Enterococcus* semi-quantitation using a single visual readout

Having confirmed visual readout of *Enterococcus* bacteria using LAMP-OSD, we sought to introduce signal thresholds within the reaction such that presence or absence of endpoint OSD fluorescence would be reflective of whether *Enterococcus* amounts were above or below the set threshold. We have previously achieved such semi-quantitation of starting template copies from a single endpoint visual readout of LAMP-OSD by integrating a false target (FT) in the reaction (**Figure 2A**).^19^ The false target is identical to the LAMP true target (TT) except the region recognized by the OSD reporter is scrambled. Consequently, it competes with the true target for amplification resources without generating a commensurate OSD fluorescence response thereby setting up a copy number threshold that the true target must exceed to generate a visible signal.

**Figure 2.**
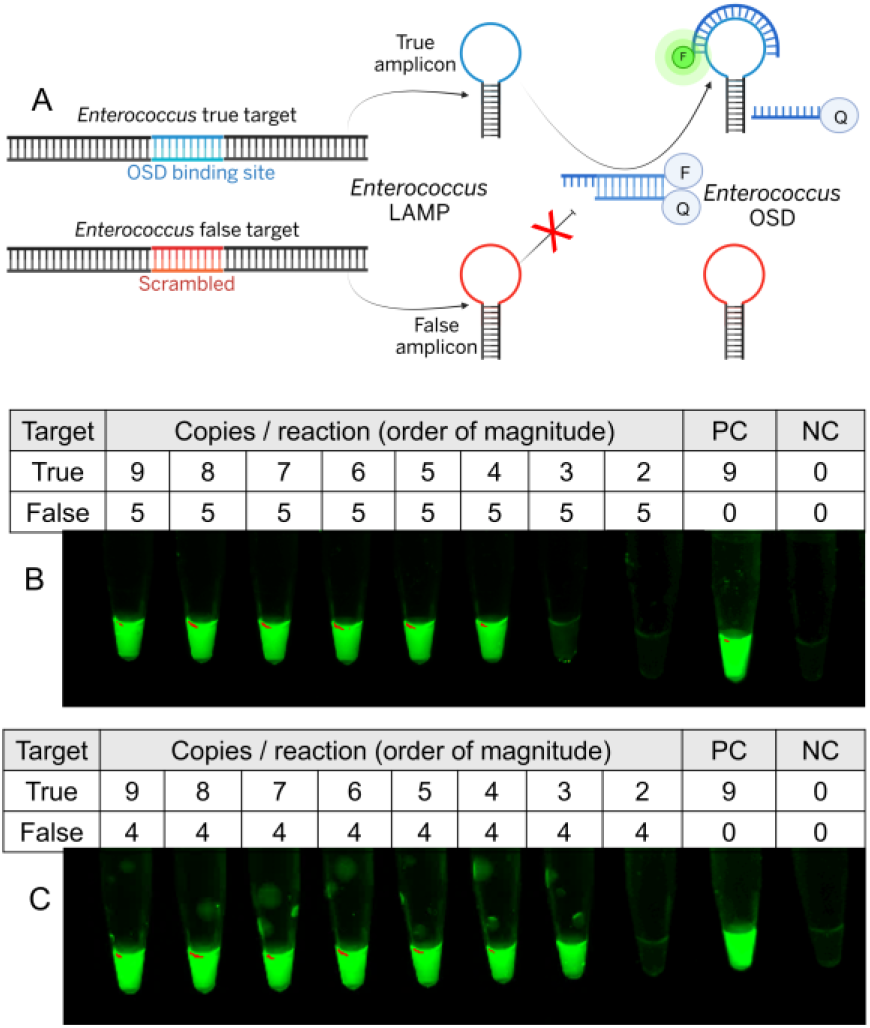
Signal thresholded LAMP-OSD analysis of synthetic *Enterococcus* DNA. (A) Schematic depicting signal thresholded LAMP-OSD. F and Q represent fluorophore and quencher, respectively. Created with BioRender.com. (B and C) Images of endpoint OSD fluorescence in a panel of thresholded *Enterococcus* LAMP-OSD assays containing indicated amounts of true and false targets. Results representative of at least triplicate experiments are depicted.

To test the application of similar signal thresholding in the *Enterococcus* assay a synthetic false target corresponding to the true target in the *Enterococcus* 23S rRNA gene was introduced into LAMP-OSD tests at about 10^5^ (**Figure 2B)** or 10^4^ (**Figure 2C**) copies/reaction. These reactions were then seeded with true templates increasing on an order of magnitude scale and observed for visual OSD fluorescence at amplification endpoint. In LAMP-OSD reactions containing on the order of 10^5^ false target copies, bright OSD fluorescence was observed only in tubes containing at least 10^4^ or more copies of true targets whereas tubes containing fewer true targets remained dark. Meanwhile, LAMP-OSD assays thresholded with ten-fold lower amounts of false targets (on the order of 10^4^ copies/reaction) produced bright OSD fluorescence when amplifying at least 10^3^ or more copies of true targets. In both cases the highest ratio of false targets / true targets that allowed bright OSD signal (inflection ratio = max(FT/TT)) was found to be about 10 (**Figure 2** and **Supplementary Figure 6**). These results suggest that false targets added to the *Enterococcus* LAMP-OSD assay predictably create readout thresholds for true target copy numbers on an order of magnitude scale. Consequently, known amounts of false targets and the inflection ratio of the thresholded LAMP-OSD assay should allow derivation of true target copies in an unknown sample.

To verify that signal thresholding *Enterococcus* LAMP-OSD assays could also be used for semi-quantitative bacterial analysis, several replicate assays containing on the order of 10^2^ or 10^4^ CFU / reaction of *E. faecalis* were spiked with false targets increasing on an order of magnitude scale. At amplification end-point, assays containing about 10^2^ CFU of *E. faecalis* produced bright OSD fluorescence only in reactions where false target thresholds were ≤10^5^ DNA copies while remaining dark in the presence of 10^6^ or more copies of false targets (**Figure 3A**). Meanwhile, 10^4^ CFU of *E. faecalis* produced bright fluorescence only when the false target threshold was set at ≤10^7^ copies. (**Figure 3B**). These results suggest that different amounts of *Enterococcus* bacteria can be distinguished on an order of magnitude scale with only a single endpoint visual readout of presence or absence of OSD signal in LAMP-OSD tests thresholded with increasing copies of false targets. The inflection ratio (max(FT/CFU)) for thresholded LAMP-OSD analysis of *Enterococcus* colony forming units was determined to be about 10^3^ (**Figure 3** and **Supplementary Figure 7**).

**Figure 3.**
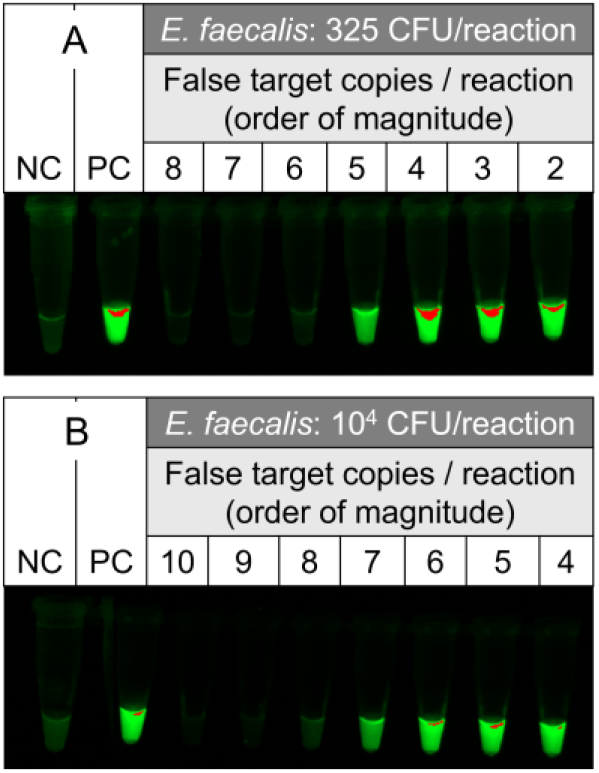
Signal thresholded LAMP-OSD analysis of *Entero-coccus faecalis* bacteria. (A and B) Images of endpoint OSD fluorescence in panels of thresholded LAMP-OSD assays containing indicated colony forming units of lab-cultivated *E. faecalis* and log10 copies/reaction of false targets. NC: negative control lacking any templates; PC: positive control containing synthetic true target templates.

### Application of thresholded LAMP-OSD for semi-quantitation of *Enterococcus* in environmental samples

Having demonstrated the ability of thresholded LAMP-OSD to distinguish lab-cultivated *Enterococcus* amounts on the log_10_ scale, we hypothesized that the thresholded LAMP-OSD assay would be able to distinguish between different levels of *Enterococcus* contamination in environmental water. To evaluate this hypothesis, 400 mL of freshwater samples were collected on different days from a local creek (**Supplementary Table 2**). Microbial contaminants in the samples were concentrated on filter paper discs and heat-lysed in 500 μL buffer prior to analysis of 2.5 μl aliquots by TaqMan qPCR and thresholded LAMP-OSD tests. *Enterococcus* colony forming units were also measured by cultivating microbial contaminants filtered from 5 mL of water on mEI agar plates.

Water samples in which *Enterococcus* could not be detected by TaqMan qPCR also failed to generate any OSD fluorescence in thresholded LAMP-OSD assays (**Figure 4A**). Water samples in which qPCR yielded a measurable average Cq (quantification cycle) of 30.23 (62 templates/reaction) but zero CFU/ml (MF) produced bright endpoint OSD fluorescence in LAMP assays thresholded with 10^3^ order of magnitude of false target copies. Assays thresholded with higher amounts of false targets remained as dark as the negative control lacking specific templates (**Figure 4B**). Water samples containing about two orders of magnitude higher levels of *Enterococcus* contamination with average qPCR Cq of 23.9 (4036 templates/reaction) and 59 CFU/ml, produced observable endpoint OSD fluorescence in LAMP assays thresholded with as much as 10^5^ orders of magnitude of false target copies (**Figure 4C**). These results demonstrate that LAMP-OSD tests performed on par with qPCR in identifying *Enterococcus-*contaminated environmental water samples. Moreover, the magnitude of the false target threshold that allowed visible endpoint OSD fluorescence was commensurate with the amount of *Enterococci* in the samples. Consequently, with a single endpoint visual assay readout the thresholded LAMP-OSD test could semi-quantitatively indicate the level of *Enterococcus* contamination in the samples – water samples with lower Enterococcal loads produced visible OSD fluorescence only at low false target thresholds while more heavily contaminated samples appeared bright at higher thresholds.

**Figure 4.**
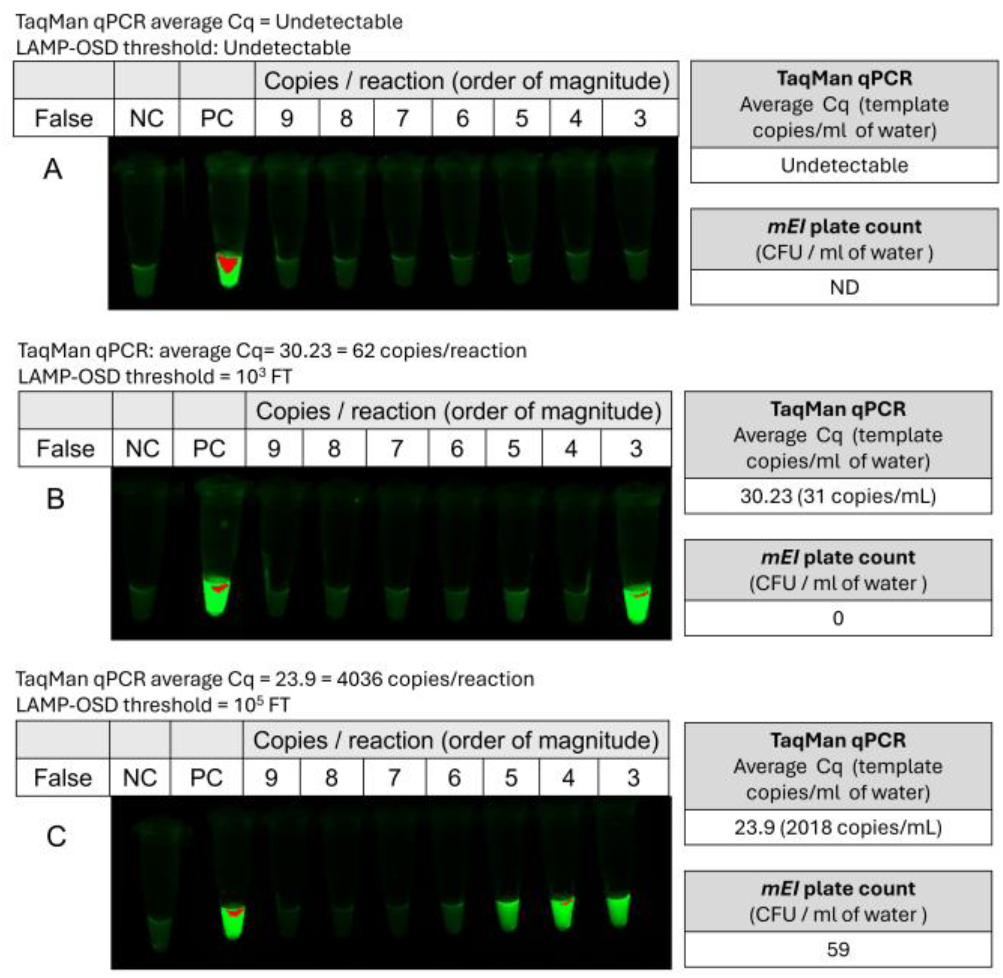
Signal thresholded LAMP-OSD analysis of *Enterococcus* bacteria in environmental freshwater samples. Images of endpoint OSD fluorescence in a panel of thresholded *Enterococcus* LAMP-OSD assays of three different water samples (A, B, and C) collected from Waller creek on different days are depicted. The OSD signal in the LAMP assays was thresholded with indicated log_10_ copies/reaction of false targets. NC: negative control lacking any templates; PC: positive control containing synthetic true target templates. Cq values were converted to copies/reaction and copies/ml of original water sample by using a standard curve (**Supplementary Figure 8**). LAMP-OSD threshold was calculated as the maximum FT amount that allowed bright OSD fluorescence (**Supplementary Figure 8**). ND: not determined.

## CONCLUSIONS

We have demonstrated a thresholded LAMP-OSD assay for semi-quantitatively assessing the level of *Enterococcus* contamination in environmental water. This assay performed on par with qPCR and was able to readily distinguish water samples with different levels of Enterococcal contamination without any interference from non-specific signal. This assay setup is much more amenable for remote field use and ‘real-time’ water quality monitoring when compared to qPCR. The thresholded LAMP-OSD assay needs only one hour of incubation at a single temperature. Furthermore, semi-quantitative test readout is accomplished with a simple visual inspection of presence or absence of OSD fluorescence at assay endpoint. By eliminating the need for thermal cycling or continuous monitoring of reaction kinetics, LAMP-OSD requires much less complex instrumentation than qPCR.

Using only a heat block and a simple visual test readout LAMP-OSD can go beyond a yes/no answer and produce a semi-quantitative estimation of the level of *Enterococci* in a sample using a panel of reactions. The simplicity of the thresholded LAMP-OSD system makes it attractive for in-field and low resource implementation, especially for widespread, frequent screening and tactical surveillance. Accessibility to on- or near-site implementation of LAMP-OSD can be further enhanced by using freeze-dried reaction mixes and low-cost remote chemical or battery-operated heaters.^11, 28, 29^ Operating costs may be further reduced by using robust engineered LAMP enzymes and low-cost production platforms^30-33^ and LAMP reactions optimized for lower temperature and therefore instrumentation needs.^34^ Subjective assessment of test results by the experimenter has proved reliable including in self-testing and clinical diagnostics.^35^ Nevertheless, readily available field-ready instruments may be used to evaluate OSD fluorescence to minimize potential user error.^36^

In future studies, large scale performance metrics and generalizability of the thresholded *Enterococcus* LAMP-OSD assay along with our previously reported assays for *E. coli* and the human fecal indicator HF183^11^ for FIB semi-quantitation will be evaluated in field studies of larger numbers, sources, and diversity of environmental and drinking water samples.

## ASSOCIATED CONTENT

**Supporting Information. Supplementary Table 1**. Oligonucleotide and template sequences used in the study. **Supplementary Figure 1**. *Enterococcus faecalis* template sequence annotated with LAMP and qPCR primers and probes. **Supplementary Figure 2**. 6-primer LAMP analysis of *Enterococcus* synthetic DNA templates. **Supplementary Figure 3**. Visual LAMP-OSD analysis of *Enterococcus* synthetic DNA. **Supplementary Figure 4**. 5-primer LAMP-OSD analysis of *Enterococcus* synthetic DNA. **Supplementary Figure 5**. LAMP-OSD analysis of *Escherichia coli* before and after transformation with an *Enterococcus* 23S rRNA encoding plasmid. **Supplementary Figure 6**. Replicates of signal thresholded LAMP-OSD analysis of synthetic *Enterococcus* DNA. **Supplementary Figure 7**. Replicates of signal thresholded LAMP-OSD analysis of *Enterococcus faecalis* bacteria. **Supplementary Table 2**. Waller Creek sample collection conditions. **Supplementary Figure 8**. Parameters for estimation of template amounts by TaqMan qPCR and thresholded LAMP-OSD. This material is available free of charge via the Internet at http://pubs.acs.org.

## AUTHOR INFORMATION

### Author Contributions

The manuscript was written through contributions of all authors. All authors have given approval to the final version of the manuscript.

## ACKNOWLEDGMENT

We would like to acknowledge support from the community of students that make up the DIY Diagnostic FRI Research Lab (https://diystream.cns.utexas.edu/).).

## ABBREVIATIONS

FIB: fecal indicator bacteria
MF: membrane filtration
MPN: most probable number
PCR: Polymerase Chain Reaction
LAMP: loop-mediated isothermal amplification
OSD: oligonucleotide strand displacement
CCE: calibrator cell equivalents
qPCR: quantitative polymerase chain reaction
CFU: colony forming units
mEI: membrane *Enterococcus* Indoxyl-β-D-Glucoside
NC: negative control
PC: positive control
TT: true template
FT: false template
SNP: single nucleotide polymorphism
DNA: deoxyribonucleic acid
ATCC: American Type Culture Collection
rpm: revolutions per minute
US EPA: United States Environmental Protection Agency.

