## Supplementary information for "Fieldable isothermal nucleic acid test for rapid semi-quantitative visual readout of *Enterococci* in recreational waters"

Supplementary Table 1
Supplementary Figure 1
Supplementary Figure 2
Supplementary Figure 3
Supplementary Figure 4
Supplementary Figure 5
Supplementary Figure 6
Supplementary Figure 7
Supplementary Table 2
Supplementary Figure 8

**Supplementary Table 1**. Oligonucleotide and template sequences used in the study.

| **Name** | **Sequence 5’ → 3’** | **References** |
|---|---|---|
| qPCR fw primer | GAGAAATTCCAAACGAACTTG | USEPA[a] |
| qPCR rev primer | CAGTGCTCTACCTCCATCATT | USEPA |
| qPCR TaqMan probe | /56-FAM/TGGTTCTCT/ZEN/CCGAAATAGCTTTAGGGCTA/3IABkFQ/ | USEPA |
| ENT-LAMP FIP (F1c-F2) | AACGTACGTGGGTTCGGTCCTTCTACCCATGTCCAGGTTGA | Martzy[b] |
| ENT-LAMP BIP (B1-B2c) | GATGAGGTGTGGGTAGCGGAGACGAGGCTAGCCCTAAAGCT | Martzy |
| ENT-LAMP F3 | CGTAGACCCGAAACCATGTG | Martzy |
| ENT-LAMP B3 | ACAGTGCTCTACCTCCATCA | Martzy |
| ENT-LAMP LoopF | GTGCGTTTTACCGCACCT | Martzy |
| ENT-LAMP LoopB | AATTCCAAACGAACTTGGAGATAGC | Martzy |
| ENT-LAMP-OSD-FAM | /56-FAM/CGCAATTCCAAACGAACTTGGAGATAGCTGGTTCTCTCCG/3InvdT/ | This study |
| ENT-LAMP-OSD-Quencher | AGCTATCTCCAAGTTCGTTTGGAATTGCG/3IABkFQ/ | This study |
| *E. faecalis* 23S rDNA LAMP template | GGCCCCTAGTCCAAACAGTGCTCTACCTCCATCATTCTCAATTCCGAGGCT AGCCCTAAAGCTATTTCGGAGAGAACCAGCTATCTCCAAGTTCGTTTGGAA TTTCTCCGCTACCCACACCTCATCCCCGCACTTTTCAACGTACGTGGGTTC GGTCCTCCAGTGCGTTTTACCGCACCTTCAACCTGGACATGGGTAGATCAC ATGGTTTCGGGTCTACGACTACATACTTATTCGCCCTATTCAGACTC | Martzy/EPA |
| False Template (Scrambled OSD binding region) | GGCCCCTAGTCCAAACAGTGCTCTACCTCCATCATTCTCAATTCCGAGGCT AGCCCTAAAGCTATTTCATCCTACTTGTCCTGGTATGAGCCAGTTGAAAGGA ATCTCCGCTACCCACACCTCATCCCCGCACTTTTCAACGTACGTGGGTTCG GTCCTCCAGTGCGTTTTACCGCACCTTCAACCTGGACATGGGTAGATCACA TGGTTTCGGGTCTACGACTACATACTTATTCGCCCTATTCAGACTC | This study |

[a]Method 1611: Enterococci in Water by TaqMan® Quantitative Polymerase Chain Reaction (qPCR) Assay. Available at https://www.epa.gov/sites/default/files/2015-08/documents/method_1611_2012.pdf.

[b]Martzy, R.; Kolm, C.; Brunner, K.; Mach, R. L.; Krska, R.; Šinkovec, H.; Sommer, R.; Farnleitner, A. H.; Reischer, G. H. A loop-mediated isothermal amplification (LAMP) assay for the rapid detection of Enterococcus spp. in water. *Water Res.* **2017**, *122,* 62-69. DOI: https://doi.org/10.1016/j.watres.2017.05.023.

/56-FAM/: 5’-end fluorescein; /3InvdT/: 3’-end inverted dT; /3IABkFQ/: 3’-end Iowa Black FQ quencher; /ZEN/: ZEN internal quencher.

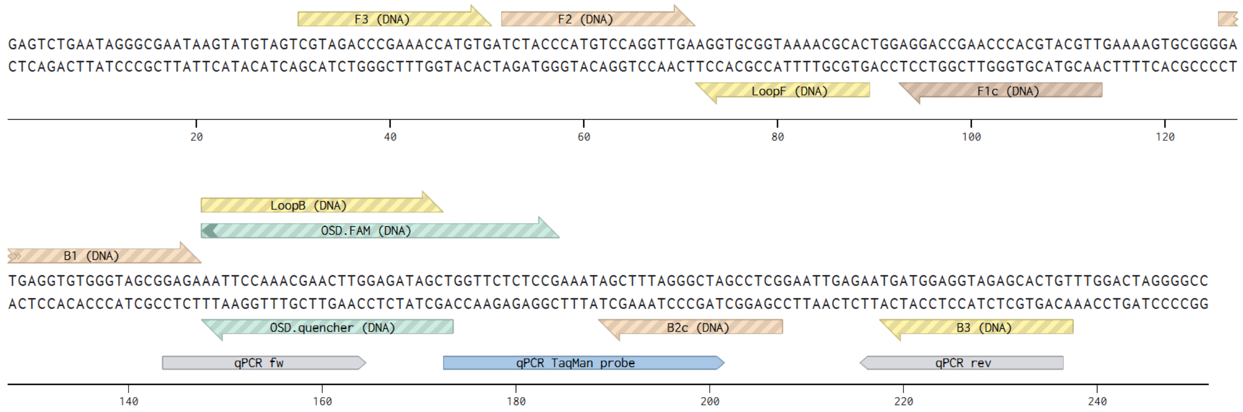


**Supplementary Figure 1**. *Enterococcus faecalis* template sequence annotated with LAMP and qPCR primers and probes.

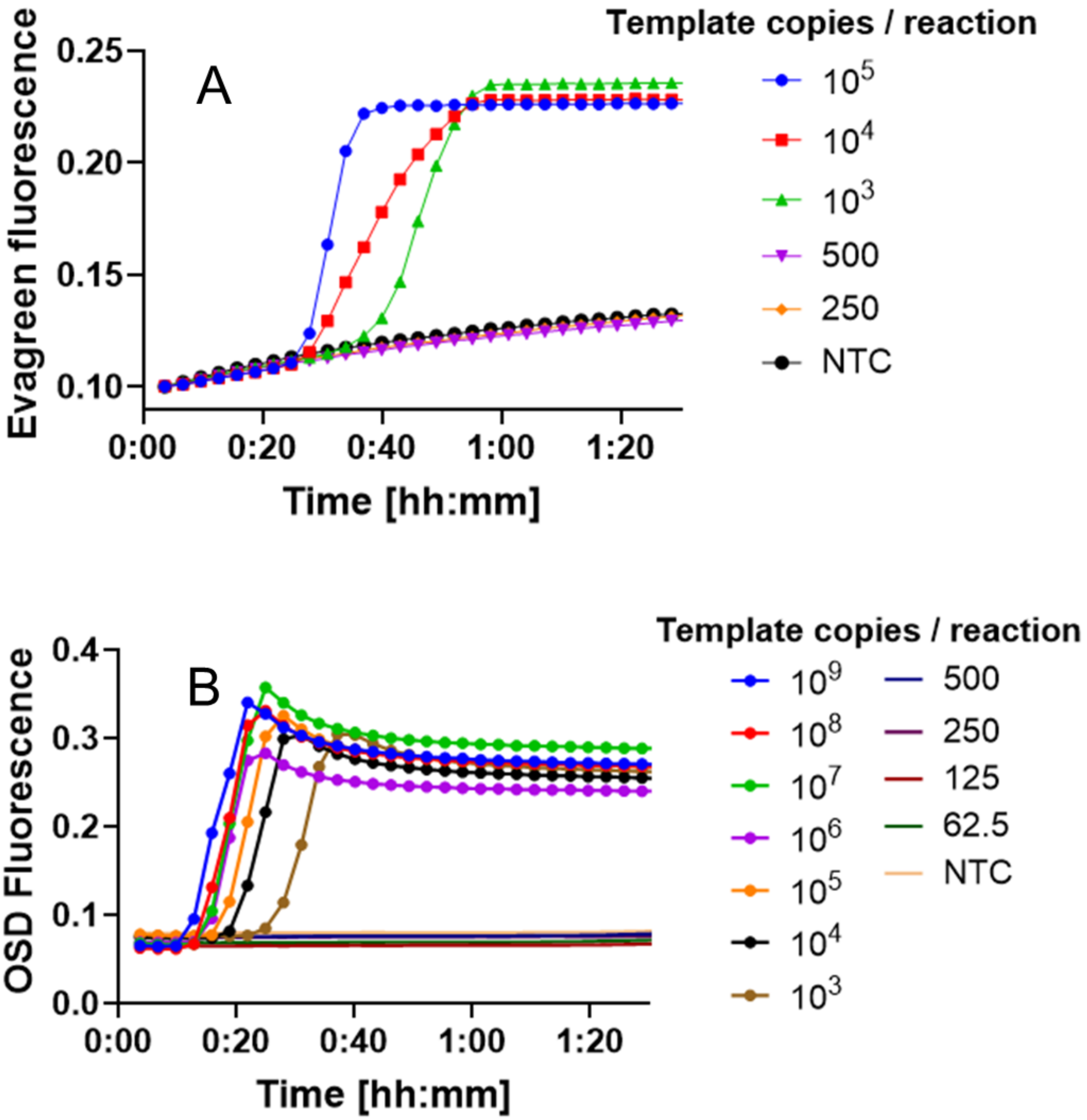


**Supplementary Figure 2. 6-primer LAMP analysis of *Enterococcus* synthetic DNA templates.** Real-time measurement of LAMP amplicon accumulation using either Evagreen dye (A) or OSD probes (B) in reactions containing indicated copies of synthetic DNA templates is depicted. Traces of reactions with detectable LAMP-OSD signals are indicated with closed circles. OSD probes were comprised of 1:5 ratio of the fluorophore and quencher labeled strands. NTC: no template control. Representative results of at least triplicate experiments are depicted.

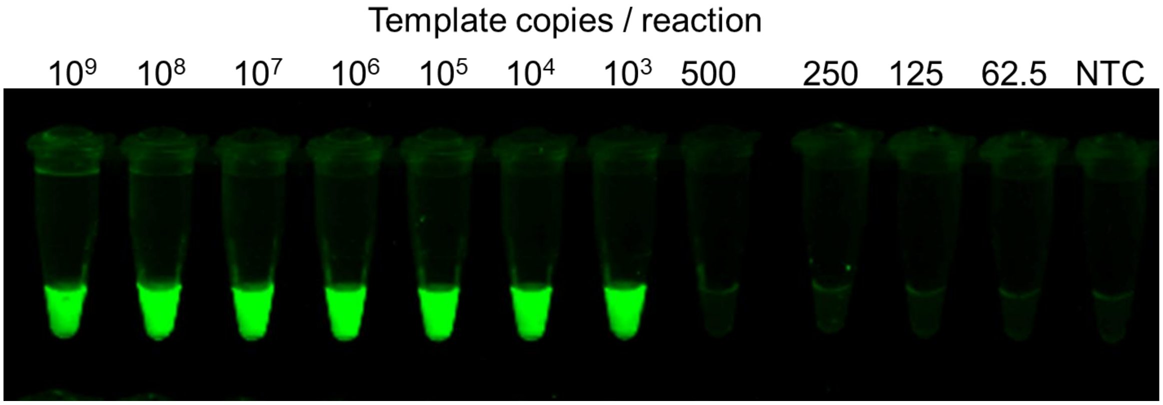


**Supplementary Figure 3. Visual LAMP-OSD analysis of *Enterococcus* synthetic DNA.** Image of endpoint OSD fluorescence in 6-primer LAMP-OSD reactions that were seeded with indicated copies of synthetic DNA templates and incubated at 65 °C for 1 hour**.** Representative results of at least triplicate experiments are depicted.

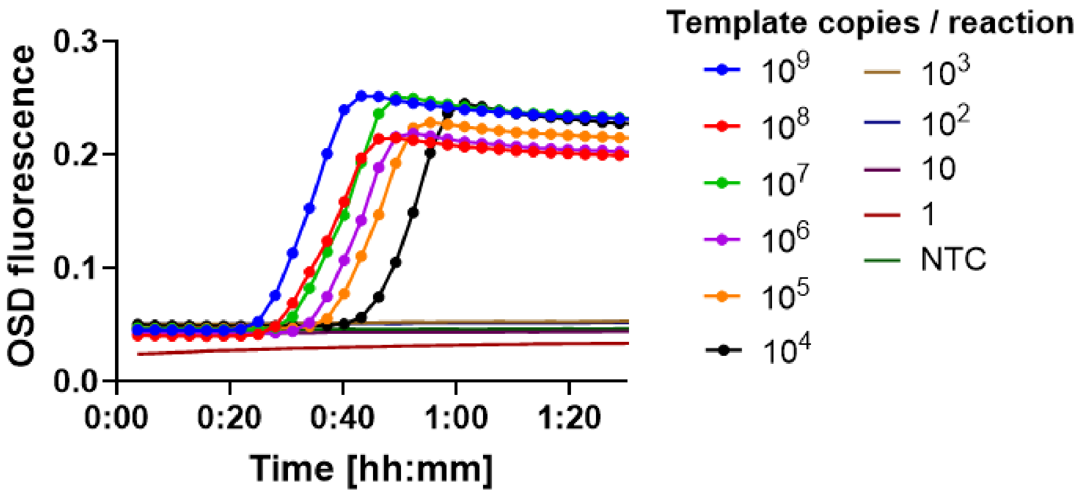


**Supplementary Figure 4. 5-primer LAMP-OSD analysis of *Enterococcus* synthetic DNA.** Real-time measurement of amplicon accumulation in 5-primer LAMP-OSD reactions lacking the loop B primers and containing indicated copies of synthetic DNA templates is depicted. Traces of reactions with detectable LAMP-OSD signals are indicated with closed circles. OSD probes were comprised of 1:1.2 ratio of the fluorophore and quencher labeled strands. NTC: no template control. Representative results of at least triplicate experiments are depicted.

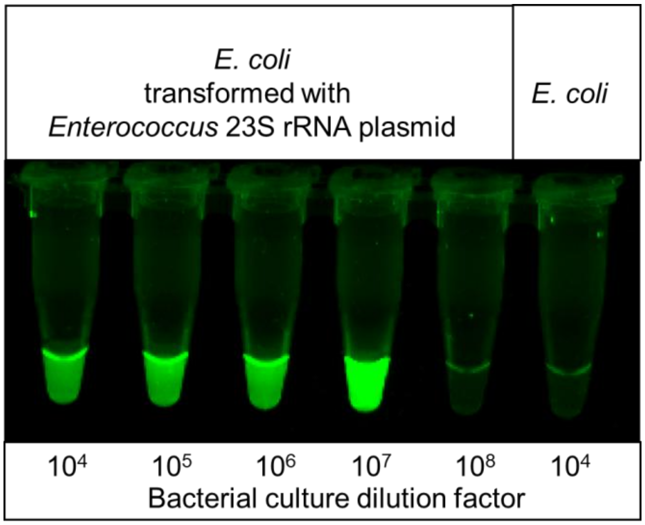


**Supplementary Figure 5. LAMP-OSD analysis of *Escherichia coli* before and after transformation with an *Enterococcus* 23S rRNA encoding plasmid.** Image of endpoint OSD fluorescence in 6-primer LAMP-OSD reactions that were seeded with indicated dilutions of logarithm phase cultures of *E. coli* bacteria that were either untransformed or were expressing a plasmid encoding the *Enterococcus* 23S rRNA sequence.

| Target | Copies / reaction (order of magnitude) | | | | | | | | PC | NC |
|---|---|---|---|---|---|---|---|---|---|---|
| True | 9 | 8 | 7 | 6 | 5 | 4 | 3 | 2 | 9 | 0 |
| False | 4 | 4 | 4 | 4 | 4 | 4 | 4 | 4 | 0 | 0 |

A

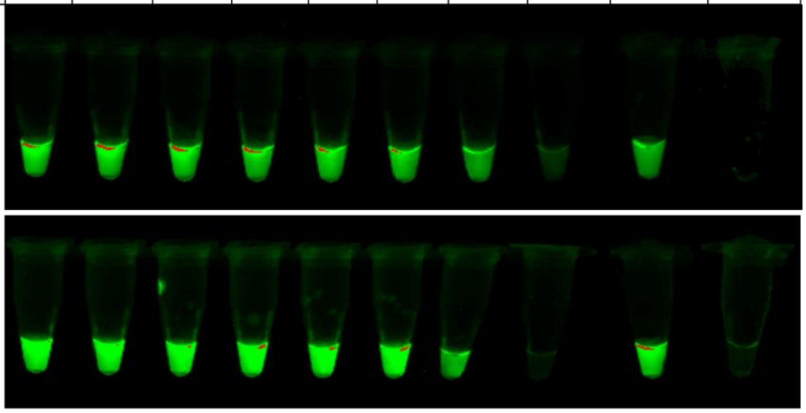

| Target | Copies / reaction (order of magnitude) | | | | | | | | PC | NC |
|---|---|---|---|---|---|---|---|---|---|---|
| True | 9 | 8 | 7 | 6 | 5 | 4 | 3 | 2 | 9 | 0 |
| False | 5 | 5 | 5 | 5 | 5 | 5 | 5 | 5 | 0 | 0 |

B

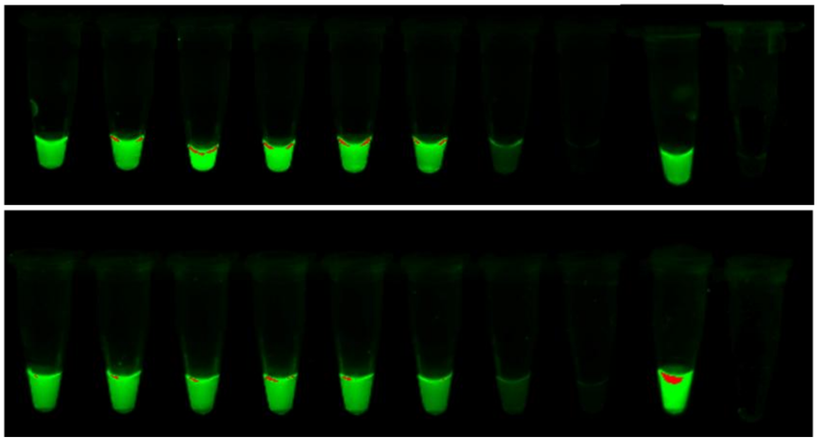

**Supplementary Figure 6. Replicates of signal thresholded LAMP-OSD analysis of synthetic *Enterococcus* DNA.** Images of endpoint OSD fluorescence in panels of thresholded *Enterococcus* LAMP-OSD assays containing indicated amounts of true *Enterococcus* and competitive false targets.

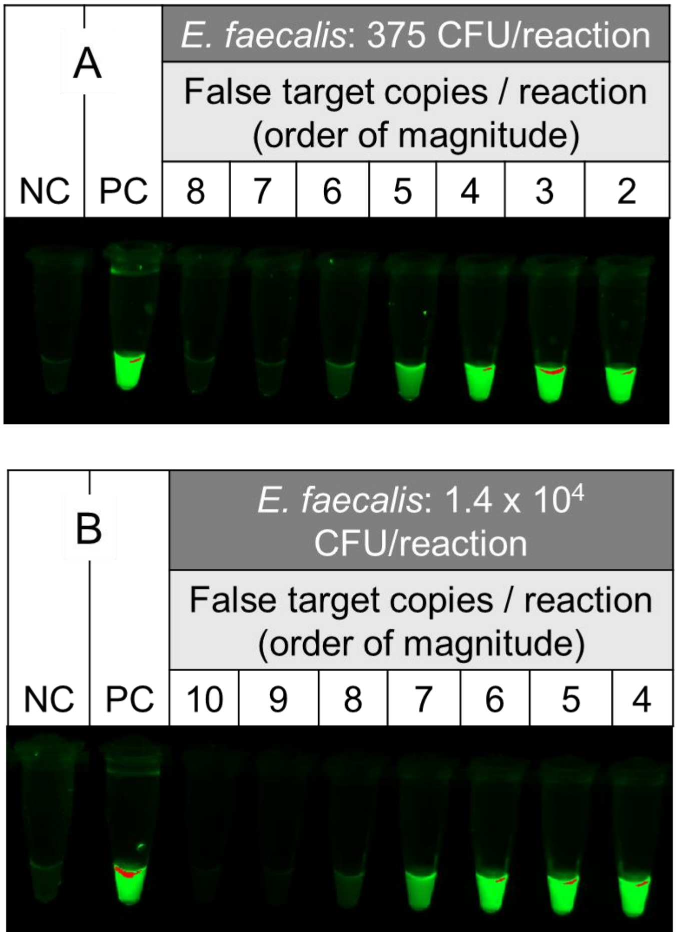


**Supplementary Figure 7**. **Replicates of signal thresholded LAMP-OSD analysis of *Enterococcus faecalis* bacteria**. Images of endpoint OSD fluorescence in panels of thresholded LAMP-OSD assays containing indicated colony forming units of lab-cultivated *E. faecalis* and log10 copies/reaction of false targets. NC: negative control; PC: positive control.

**Supplementary Table 2. Waller Creek sample collection conditions.**

| Date | *Enterococcus* levels | Location | Conditions during sampling | | | | | Notes |
|---|---|---|---|---|---|---|---|---|
| | | | Time | Air temperature | Humidity | Wind | Amount of trash | |
| 10/2/2024 | Not detected | Creek Side Hall | 2:00 PM | 32 °C | 34% | 5 mph | Light, wrappers | No rains for several weeks |
| 11/6/2024 | High | Creek Side Hall | 9:30 AM | 18 °C | 61% | None | Light, wrappers & some plastic | Rain on 11/4 and 11/5, noted rushing water at sampling site |
| 1/23/2025 | Low | Creek Side Hall | 3:30 PM | 12 °C | 21% | 7 mph | Light, wrappers & a can | Light snow two days prior to sampling |

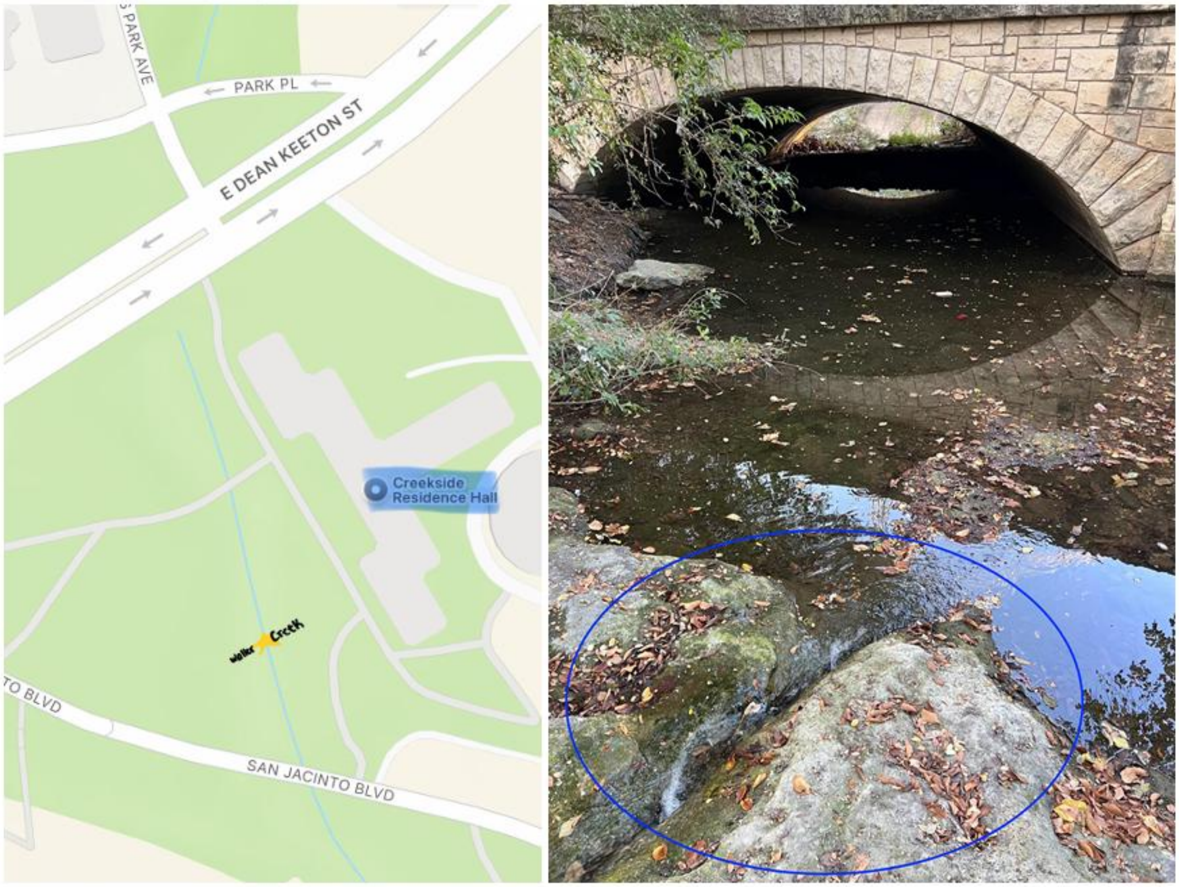


**Location map and image of Creek Side Hall sampling site.**

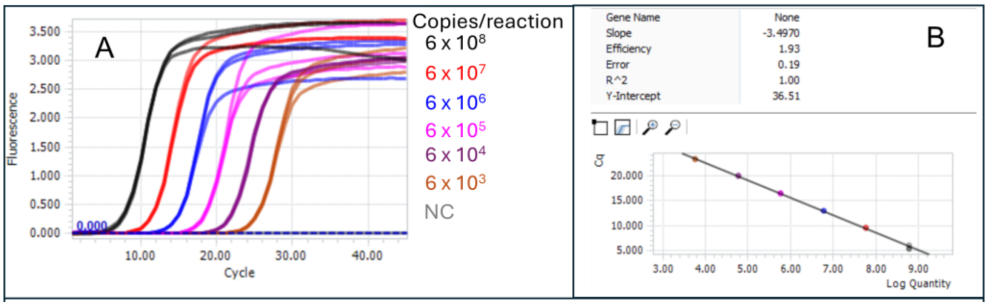


C

$$Cq = -3.4970 \times Log\ Quantity + 36.51$$

$$Copies\ per\ ml\ of\ water\ sample = \frac{\frac{Copies\ per\ reaction}{Sample\ volume\ per\ reaction} \times \text{Concentrated sample volume}}{Original\ sample\ volume}$$

$$Sample\ volume\ per\ reaction = 2.5\ \mu l$$

$$Concentrated\ sample\ volume = 500\ \mu l$$

$$Original\ sample\ volume = 400\ ml$$

D

$$LAMP - OSD\ threshold = highest\ FT\ amount\ that\ allows\ bright\ OSD\ fluorescence\ from\ TT$$

**Supplementary Figure 8**. **Parameters for estimation of template amounts by TaqMan qPCR and thresholded LAMP-OSD.** (A) *Enterococcus* TaqMan qPCR standard curve. Triplicate amplification kinetics of indicated copies of synthetic *Enterococcus* DNA templates. NC: negative control. (B) Standard curve analysis of amplification data using the LightCycler Abs Quant analysis. (C) Equations used for calculation of copies / reaction and copies/ml of water sample using qPCR data. (D) Metric for calculation of LAMP-OSD threshold.
